# Transcranial photobiomodulation with infrared laser increases power of brain oscillations

**DOI:** 10.1101/535757

**Authors:** Xinlong Wang, Jacek P. Dmochowski, Li Zeng, Elisa Kallioniemi, Mustafa Husain, F. Gonzalez-Lima, Hanli Liu

**Author notes:** Corresponding author: **Hanli Liu, Ph.D.**, Bioengineering Department, University of Texas at Arlington, Arlington, TX, 76019.

## Abstract

Non-invasive transcranial photobiomodulation with a 1064-nm laser (tPBM_L1064_) has been reported to improve human performance on cognitive tasks as well as locally upregulate cerebral oxygen metabolism and hemodynamics. However, it is unknown whether tPBM_L1064_ also modulates electrophysiology, and specifically neural oscillations, in the human brain. The hypothesis guiding this study was that tPBM_L1064_ of the right prefrontal cortex enhances neurophysiological rhythms at specific frequency bands in the human brain under resting conditions. To test this hypothesis, we recorded the 64-channel scalp electroencephalogram (EEG) before, during, and after the application of 11 minutes of 4-cm-diameter tPBM_L1064_ to the right forehead of human subjects (n=20) using a within-subject, sham-controlled design. Time-resolved scalp topographies of EEG power at five frequency bands were computed to examine tPBM_L1064_-induced EEG power changes across the scalp. The results showed time-dependent, significant increases of EEG spectral powers at the alpha (8-13 Hz) and beta (13-30 Hz) bands at broad scalp regions, exhibiting a front-to-back pattern. The findings provide the first sham-controlled topographic mapping that tPBM_L1064_ increases the strength of electrophysiological oscillations (alpha and beta bands), while also shedding light on the mechanisms of tPBM in the human brain.

## 1. Introduction

Photobiomodulation (PBM) is a process that uses light, with non-destructive and non-thermal delivery techniques, to modulate mitochondrial respiration and cellular functions in many cell types, including neurons (1–4). PBM has also been called Low-Level Laser (Light) Therapy (LLLT) when its goal is to serve as a therapy for a variety of medical conditions, such as for pain relief and wound healing (5, 6). When PBM is aimed at the human head, the term transcranial photobiomodulation (tPBM) has been used to emphasize that the target modulated by light is the human cerebral cortex, up to ~3 cm below the human scalp (7), with the goal of enhancing cerebral oxygenation and cognitive functioning (8–10). In general, most research studies or clinical applications with tPBM have utilized either lasers or light emitting diodes (LEDs) in the red (620-680 nm) or near infrared (800-980 nm) spectral range (11).

In recent years, a 1064-nm infrared laser has been utilized to apply tPBM and has been reported to improve cognitive performance on various tasks in sham-controlled studies (12–15). Moreover, we have recently demonstrated that tPBM by 1064-nm laser (tPBM_L1064_) induced significant changes in cerebral blood oxygenation as measured with functional near-infrared spectroscopy (fNIRS) (16, 17). These changes were shown to be preceded by concomitant neuron-photon interaction induced increases in the concentration of oxidized cytochrome-c-oxidase (CCO) (17), the enzyme at the terminal stage of the mitochondrial respiratory chain (18).

Because the term tPBM in the literature covers a broad spectral range (600-980 nm) of laser or LEDs, we here employ the term tPBM_L1064_ to refer to our specific application of tPBM with a 1064-nm laser. There have been no comparable human studies of CCO photo-oxidation using tPBM at wavelengths shorter than 1064 nm.

The previous *in vivo* hemodynamic and metabolic tPBM_L1064_ studies in humans provided insights into the physiological changes evoked by stimulation with a 1064-nm infrared laser (17, 18). The purported mechanism of tPBM_L1064_ in improving human cognitive function was a neuron-photon interaction initiated by CCO photo-oxidation (8, 17). In brief, incident photons cause CCO to lose electrons (i.e., to become photo-oxidized) since CCO is the main photon acceptor in cells. When CCO becomes oxidized, it has a conformation with more affinity to catalyze the reduction of oxygen to water in cellular respiration. The increment of oxidized CCO concentration therefore accelerates ATP production via oxidative phosphorylation by consuming more oxygen molecules (8, 17, 19). The faster consumption rate of oxygen molecules triggers an increase in blood flow and thus hemodynamic oxygenation from nearby blood vessels. The increased ATP in neurons is then utilized as an energy supply to carry out a variety of cellular activities and biochemical reactions (8, 19).

Since the most energy-demanding cellular activity in neurons involves maintaining membrane electrical potentials (20), the tPBM_L1064_-induced increase in oxidized CCO and ATP concentrations in neurons may potentially lead to electrophysiological alterations. Moreover, according to the findings in our previous studies, local hemodynamic and metabolic responses were reported as responses to tPBM_L1064_ on the right prefrontal cortex. However, it is yet unknown if the tPBM_L1064_ induces distal/global electrophysiological responses due to neuronal connectivity. To date, only a couple of *in vivo* studies have investigated the electrophysiological changes driven by tPBM_L1064_ (15, 21). The primary goal of this study was to determine whether tPBM_L1064_ modulates the amplitude of human cortical rhythms as measured by the scalp EEG before, during, and after tPBM.

The scalp EEG is a time-resolved measure of the aggregate synaptic activity across millions of cells in the underlying cerebral cortex. EEG has been often combined with brain stimulation (e.g. transcranial direct current stimulation (tDCS) and transcranial magnetic stimulation (TMS)) to study the mechanisms of neuromodulation techniques (22–24). Modulation in the activity of alpha (8-13 Hz), beta (13-30 Hz), and gamma (30-70 Hz) rhythms have long been associated with cognitive function and brain states (25–27). Moreover, a variety of forms of tPBM (using either LEDs at wavelengths between 800-850 nm or a laser at 1064 nm) have been reported to enhance human cognitive functions, such as working memory, sustained attention, category learning and executive skills (8, 10–15, 28).

To the best of our knowledge, few studies have reported EEG spectral power changes in response to tPBM_L1064_ (except for two preliminary studies from our own groups (15, 21)). Therefore, in this study we focused on two major research questions. First, does tPBM_L1064_ significantly modulate human cortical rhythms as compared to sham stimulation? If so, what frequency bands are responsive to tPBM_L1064_? Second, what is the spatial distribution of tPBM_L1064_ induced alterations in oscillatory power? Specifically, we hypothesized that tPBM_L1064_ of the right prefrontal cortex enhances neurophysiological rhythms at specific frequency bands in the human brain under resting conditions.

To test our hypothesis, we recorded the scalp EEG before, during, and after the application of 1064-nm infrared laser on the right frontal forehead in n=20 subjects using a within-subjects, sham-controlled design. By the end of this study, the hypothesis was confirmed by our observations that tPBM_L1064_ gave rise to significant increases in the spectral power strength of electrophysiological oscillations at alpha and beta bands across anterior to posterior regions in the human brain, revealing electrophysiological mechanisms of action of tPBM_L1064_ on the human brain.

## 2. Materials and Methods

### 2.1 Participants

Twenty healthy human participants with an average (± SD) age of 26.8 ±8.8 years were recruited from the local community of the University of Texas at Arlington. Specifically, 7 females (24.0±4.1 years of age) and 13 males (28.2±10.3 years of age) participated in the study; the two gender groups had no significant age difference (with a two-tailed t-test, *p*>0.2). The exclusion criteria included: (1) previous diagnosis with a psychiatric disorder, (2) history of neurological disease, (3) history of severe brain injury, (4) history of violent behavior, (5) prior institutionalization/imprisonment, (6) current intake of any psychotropic medicine or drug, (7) previous diagnosis with diabetes, (8) history of smoking, (9) excessive alcohol consumption or (10) pregnancy. The study protocol was approved by the Institutional Review Board (IRB) at the University of Texas at Arlington. Informed consent was obtained prior to all experiments.

### 2.2 Experimental setup for simultaneous EEG-tPBM_L1064_ measurements

We employed a FDA-cleared, 1064-nm, continuous-wave laser (Model CG-5000 Laser, Cell Gen Therapeutics LLC, Dallas, TX, USA) used in previous studies (12–17, 28). The laser’s aperture delivered a well-collimated beam with an area of 13.6 cm^2^. To avoid thermal sensation for a more truthful sham-controlled study, we decreased the laser power to 2.2 W from 3.4 W used in previous studies (12–17, 28). Indeed, all participants reported no thermal sensation. This laser power of 2.2 W resulted in a power density of 0.162 W/cm^2^, energy density of 9.72 J/cm^2^ per minute, and a total energy dose of 1452 J over the 11-min tPBM_L1064_ duration (2.2 W x 660 s = 1452 J). Note that the optical energy density (or fluence) received on the cortical region would be only 1-2% of that delivered on the scalp by the laser aperture, based on recent literature (7, 29–31). For sham stimulation, the laser power output was set to 0.1 W while a black cap was also placed in front of the aperture to further block the laser. The corresponding power density under the sham stimulation was further tested to be 0 mW/cm^2^ by a sensitive power meter (Model 843-R, Newport Corporation, MA) to ensure complete blockage of laser light.

The experimental setup of the simultaneous EEG-tPBM_L1064_ measurements is shown in Figs. 1(a) and 1(b). The EEG data were collected using a 64-channel EEG instrument (ActiveTwo, Biosemi, Netherlands). Channel 48, Cz, was used as the reference. tPBM_L1064_ was applied by non-contact delivery to the right forehead of each subject at a frontal site (near the Fp2 location of the international 10-10 system), the same laser light delivery location demonstrated to enhance cognitive performance and local oxygen metabolism in our previous studies (13–15, 17). Laser protection goggles with extra black-tape rims were worn by the subject and the experimental operator throughout the entire experiment, before, during, and after tPBM_L1064_. To prevent sleepiness or drowsiness, the subjects kept their eyes open during the EEG-tPBM_L1064_ measurements after the laser aperture was well positioned on their forehead. Each subject wore an EEG cap with 64 electrodes positioned according to the standard 10-10 EEG electrode placement (32).

**Figure 1.**
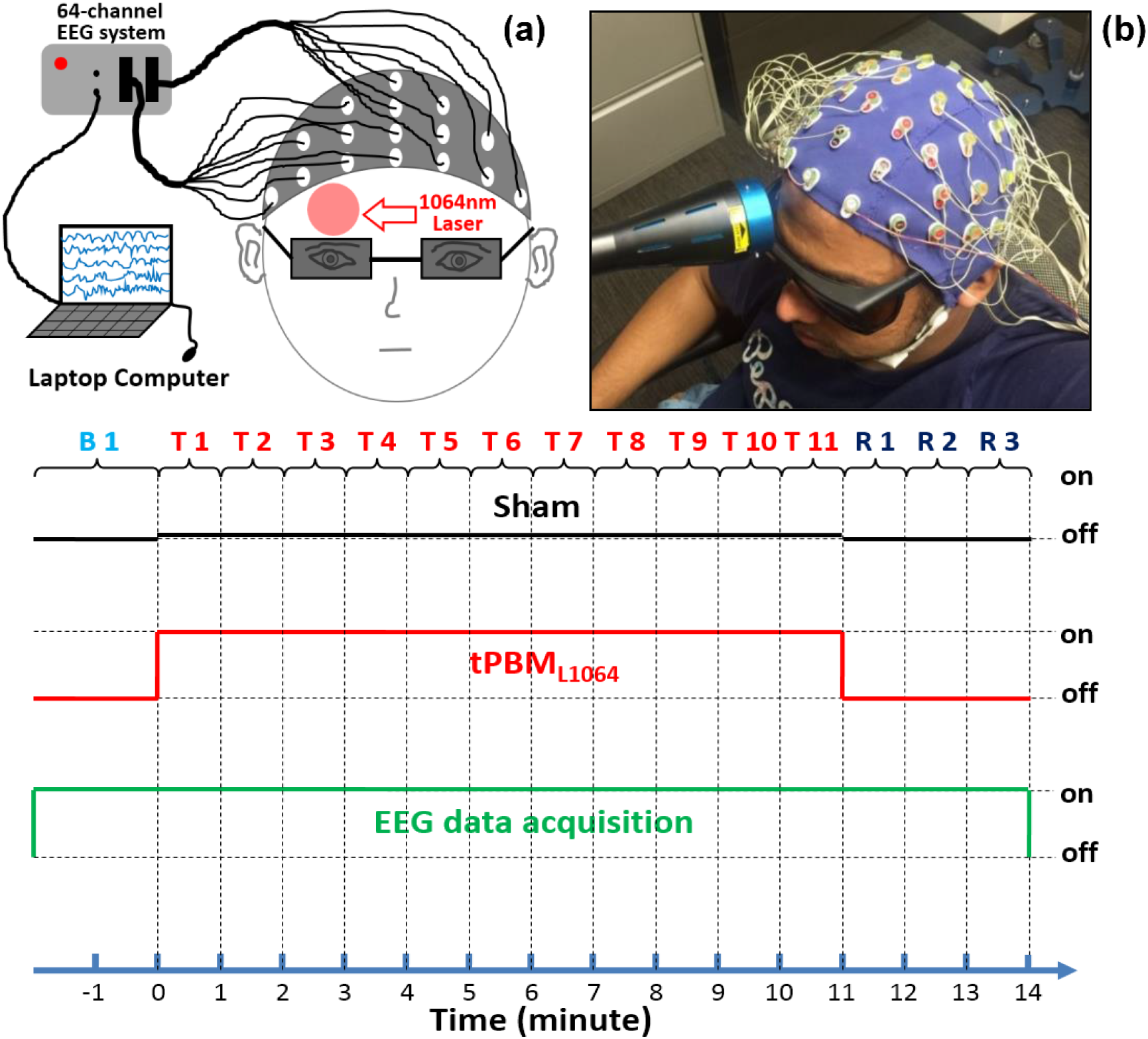
(a) A highly schematic drawing of the setup, (b) a photo of actual setup for the EEG-tPBM_L1064_ experimental setup, and (c) illustration of the study protocol for EEG-tPBM_L1064_ and EEG-Sham experiments.

### 2.3 Experiments of simultaneous EEG-tPBM_L1064_

Figure 1(c) shows the experimental design for both sham and active tPBM_L1064_. Experiments consisted of a 2-minute baseline period (Bl), an 11-minute laser stimulation period (T1-T11), and a 3-minute recovery period (R1-R3). All 64-channel EEG data were acquired throughout the 16-minute duration. Each subject received both active and sham stimulation on separate days spaced by 14 days, with the order of the sessions randomized and balanced across subjects. All subjects were unaware that one of the sessions consisted of sham stimulation. After each experiment, each subject was asked about perceptions of heat from the stimulation.

### 2.4 EEG data analysis

We conducted data analysis on 64-channel EEG time series in several steps.

#### Pre-processing

All 64-channel EEG time series were pre-processed using the EEGLAB toolbox on the Matlab platform (33). Each of the raw time series was first band-pass filtered between 1-70 Hz and then notch filtered to remove 60-Hz line noise. To remove artifacts from eye blinks, saccades, and jaw clenches, we performed Independent Component Analysis (ICA) (34) via the EEGLAB function “runica”. Artifactual components were visually identified and removed from the data (35, 36). Electrode Cz was used as the reference for the other 63 EEG channels.

#### Selection of frequency bands

For spatial topographies of EEG power spectral densities, we selected the five commonly used EEG frequency bands; namely, delta (0.5-4 Hz), theta (4-8 Hz), alpha (8-13 Hz), beta (13-30 Hz), and gamma (30-70 Hz) frequency bands.

#### Baseline quantification

For each of these five frequency bands, the 60-sec pre-laser (or baseline) EEG power spectral densities were temporally averaged for each EEG channel for both active and sham tPBM_L1064_ group data. Then, we performed a two-sample t-test between the EEG baselines during active and sham tPBM_L1064_ for each of 64 channels. The results showed no statistically significant difference between the baselines of the active and sham group data for each of the five bands.

#### Time-resolved, baseline-normalized topography

For each frequency band, we created time-dependent, 64-channel (i.e., 64×1) power density vectors, P_tPBM_ or P_sham_, for either tPBM_L1064_ or sham experiment during the stimulation (11 min) and recovery (3 min) period, as well as a corresponding baseline power density vector, P_base_ (based on the 2^nd^ minute baseline data). Baseline-normalized, tPBM_L1064_-induced (or sham-induced) increases in power density were quantified as nPtPBM_L1064_ = P_tPBM_/P_base_ (or nP_sham_ = P_sham_/P_base_) for each of the 14 time-dependent periods during and after tPBM_L1064_ (i.e., T1-T11 and R1-R3). This data processing routine is shown in Fig. 2. Next, after close inspection on the magnitudes of time-resolved, baseline-normalized EEG power density for each EEG channel at each frequency band, we excluded 3 subjects as outliers. The exclusion criterion was that any subject having laser-induced power alteration (i.e., increase or decrease) larger than 100% in magnitude would be excluded as an outlier, regardless of any channel or/and any frequency band.

**Figure 2.**
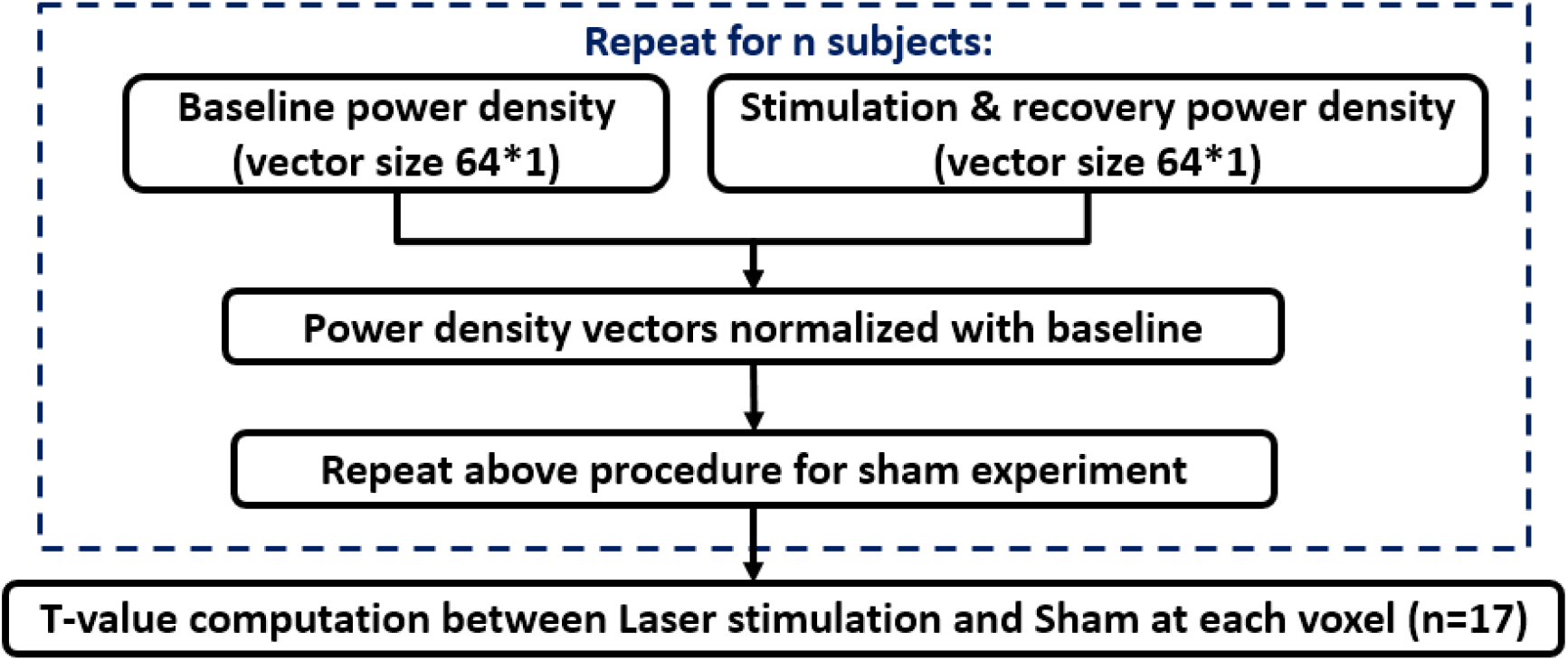
A flow chart for EEG data analysis to obtain topographic maps of EEG power spectral density differences between active and sham tPBM_L1064_. This process routine was used for each of the five frequency bands.

#### Statistical analysis

Mean values of nPtPBM_L1064_ and nP_sham_ over 17 participants at each temporal point were calculated, and then the mean differences with respect to the baseline power density were obtained and presented by 2D topographic maps in temporal sequence (in min). Next, we performed two-sample t-tests between mean nPtPBM_L1064_ and nP_sham_ (i.e., comparing active and sham tPBM_L1064_ experiments) in temporal sequence, with a statistical significance level chosen to be p<0.05 after Bonferroni correction (see Fig. 2), which is commonly used for multiple comparison to reduce Type I error. A total of 14 statistical T-maps were made, for each frequency band, to visually show regions where tPBM_L1064_ induced significant increases in EEG power densities on the human head. Moreover, using another way to present statistically significant results, we calculated time-resolved effect size for the group differences using Cohen’s “*d*” (37) and displayed it on topographical maps, following the similar presentation given in (38).

## 3. Results

A sham-controlled within-subjects design with n=20 healthy human participants was employed to assess the effect of tPBM_L1064_ on the power of EEG oscillations at four canonical bands. After excluding 3 subjects whose time-resolved magnitudes of baseline-normalized EEG power density met the exclusion criterion, we report group effects derived from the remaining 17 participants.

Figure 3(a) shows time-resolved spatial topographies of group-level differences in baseline-normalized EEG power density at the delta (0.5-4 Hz) frequency band between tPBM_L1064_ and sham experiments (i.e., nP_tPBM_ - nP_sham_) and corresponding statistical t-maps, during the 11 minutes of tPBM_L1064_/sham (T1, T2, … T11) and 3 minutes of recovery (R1, R2, and R3) period. Figure 3(b) presents the corresponding topographies for the theta (4-8 Hz) frequency band. Figure 3(a) does not show obvious increases in mean magnitude of EEG delta power density at any temporal point with respect to the sham-induced results, while Fig. 3(b) seems to reveal mean magnitude increases of tPBM_L1064_-induced EEG theta power density at several temporal periods (e.g., time=3, 8, 11 min) compared to the sham results. However, the respective T-maps after Bonferroni correction indicated that the magnitude of laser-induced changes of EEG powers in both delta and theta bands were not statistically significant across any 14 time periods, suggesting that tPBM_L1064_ had no statistical effects on delta and theta bands.

**Figure 3.**
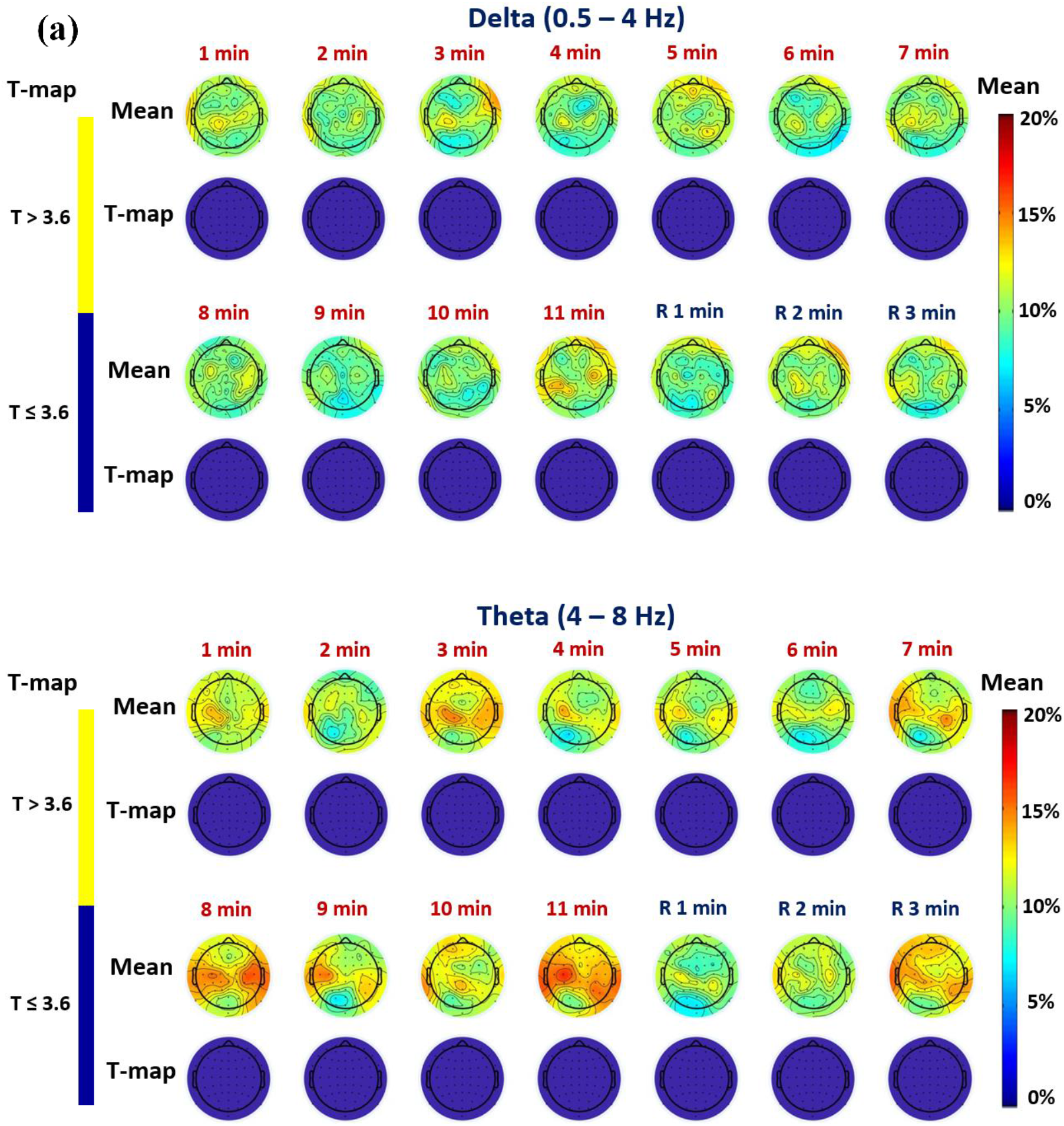
Time-resolved topographical maps of group-level differences in baseline-normalized EEG power density at (a) delta and (b) theta bands between tPBM_L1064_ and sham experiments (i.e., nP_TPBM_ - nP_sham_) and corresponding statistical T-maps, during the 11 minutes of tPBM_L1064_/sham (T1, T2, … T11) and 3 minutes of recovery (R1, R2, and R3) period. For both (a) and (b) panels, the color bar on the right side represents mean difference in percentage increase of respective power density between tPBM_L1064_ and sham stimulations. The color bar on the left side represents the T-score cutoff for the Bonferroni-corrected p-value (p < 0.05, which is equivalent to T>3.6) of significant differences in power density between tPBM_L1064_ and sham stimulations.

Figure 4(a) illustrates 14 time-resolved (i) spatial topographies of group-level mean differences in baseline-normalized EEG power density at the alpha (8-13 Hz) frequency band between tPBM_L1064_ and sham experiments, (ii) corresponding statistical T-maps, and (iii) respective effect-size topographic maps, during and after tPBM_L1064_. This figure shows that alpha power increased in mean magnitude after tPBM_L1064_ onset, and remained elevated over bilateral anterior and posterior regions even after stimulation. Statistical T-maps after Bonferroni correction showed that stimulation of the right forehead by tPBM_L1064_ increased alpha oscillation powers significantly over many bilateral regions, anteriorly and posteriorly, on the human head throughout the stimulation time, particularly 8-11 minutes after the stimulation onset. This observation was supported by the effect-size maps, which demonstrated that tPBM_L1064_ caused very large effects in alpha power as compared to the sham control in the above mentioned regions and temporal periods, given that |d| < 0.2 – small effect; 0.2 < |d| < 0.8 – medium effect; |d| > 0.8 – large effect; |d| > 1.2 – very large effect (38).

**Figure 4.**
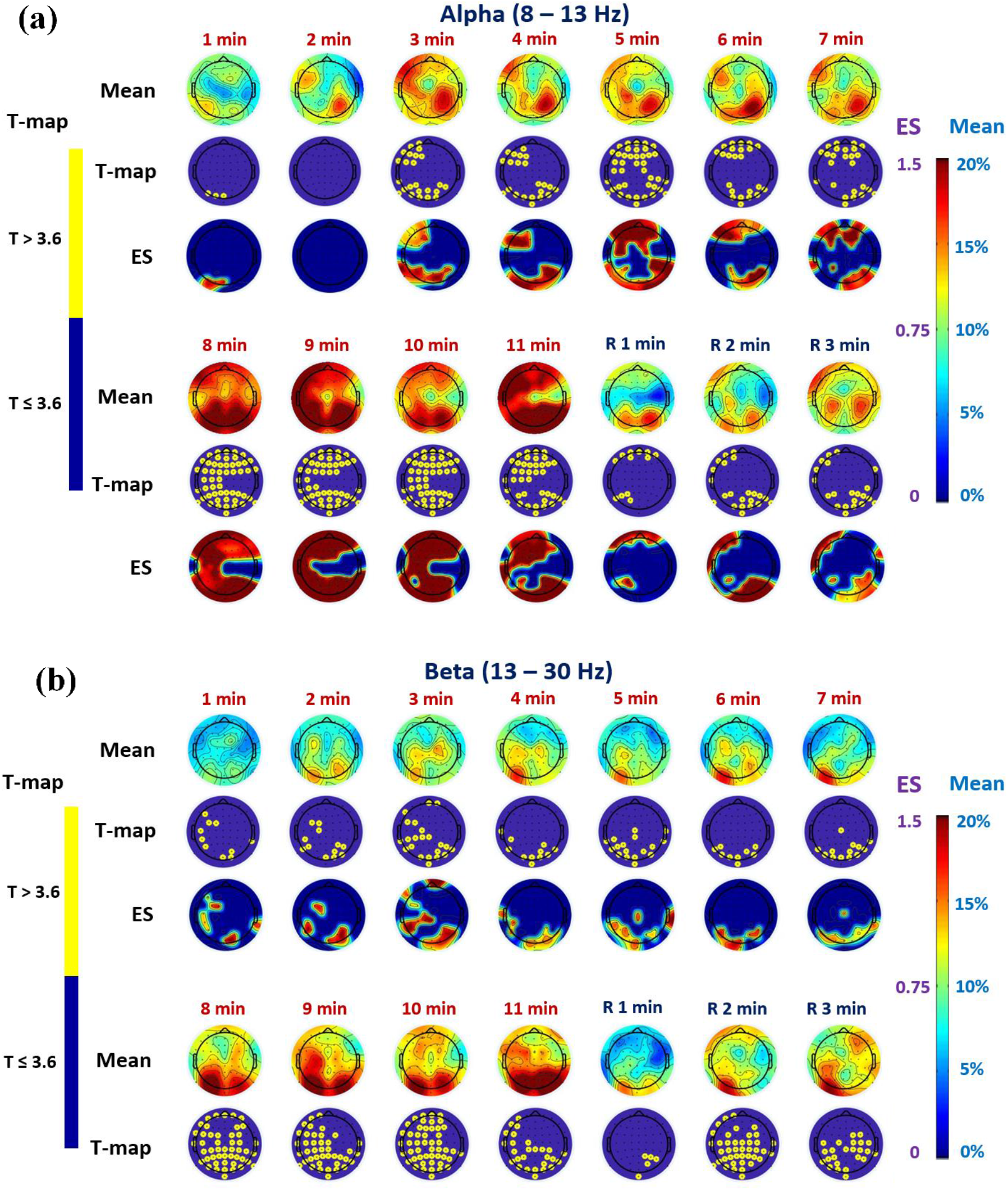
Time-resolved topographical maps of group-level differences in baseline-normalized EEG power density at (a) alpha and (b) beta band between tPBM_L1064_ and sham experiments (i.e., nP_TPBM_ - nP_sham_), corresponding statistical t-maps, and respective effect size maps, during the 11 minutes of tPBM_L1064_/sham (T1, T2, … T11) and 3 minutes of recovery (R1, R2, and R3) period. For both (a) and (b) panels, the color bar on the right side represents both effect size (ES) and mean difference in percentage increase of respective power density between tPBM_L1064_ and sham stimulations. The color bar on the left side represents the T-score cutoff for the Bonferroni-corrected p-value (p < 0.05, which is equivalent to T>3.6) of significant differences in power density between tPBM_L1064_ and sham stimulations.

Figure 4(b) illustrates 14 time-resolved (i) spatial topographies of group-level mean differences in baseline-normalized EEG power density at the beta (13-30 Hz) frequency band between tPBM_L1064_ and sham experiments, (ii) corresponding statistical T-maps, and (iii) respective effect-size topographic maps, during and after tPBM_L1064_. Increases of beta-band power were also observed, occurring mainly over the posterior electrodes 4-7 minutes after stimulation onset. Similar to the alpha-band, tPBM_L1064_-induced beta oscillation powers were elevated significantly across most of the scalp (both anterior and posterior) 8-11 minutes after tPBM_L1064_ onset. The statistical T-maps shown in Fig. 4(b) also revealed statistically significant differences in beta power density between active and sham tPBM_L1064_ during the recovery period (i.e., during R1, R2, and R3) in several scalp regions. This observation was further confirmed by the respective effect-size maps, illustrating that tPBM_L1064_ caused very large effects in beta power as compared to the sham control in the above mentioned regions and temporal periods, given that |d| < 0.2 – small effect; 0.2 < |d| < 0.8 – medium effect; |d| > 0.8 – large effect; |d| > 1.2 – very large effect (38).

Regarding electrophysiological responses of the brain to tPBM_L1064_ at the gamma (30-70 Hz) frequency band, Fig. 5 seems to show some mean magnitude changes anteriorly and posteriorly, particularly during 8-11 min after the stimulation onset. However, the Bonferroni-corrected statistical analysis as shown by corresponding T-maps in Fig. 5 indicates that the laser-induced changes of EEG gamma powers were not statistically significant between the laser and sham stimulation across any 14 time periods.

**Figure 5.**
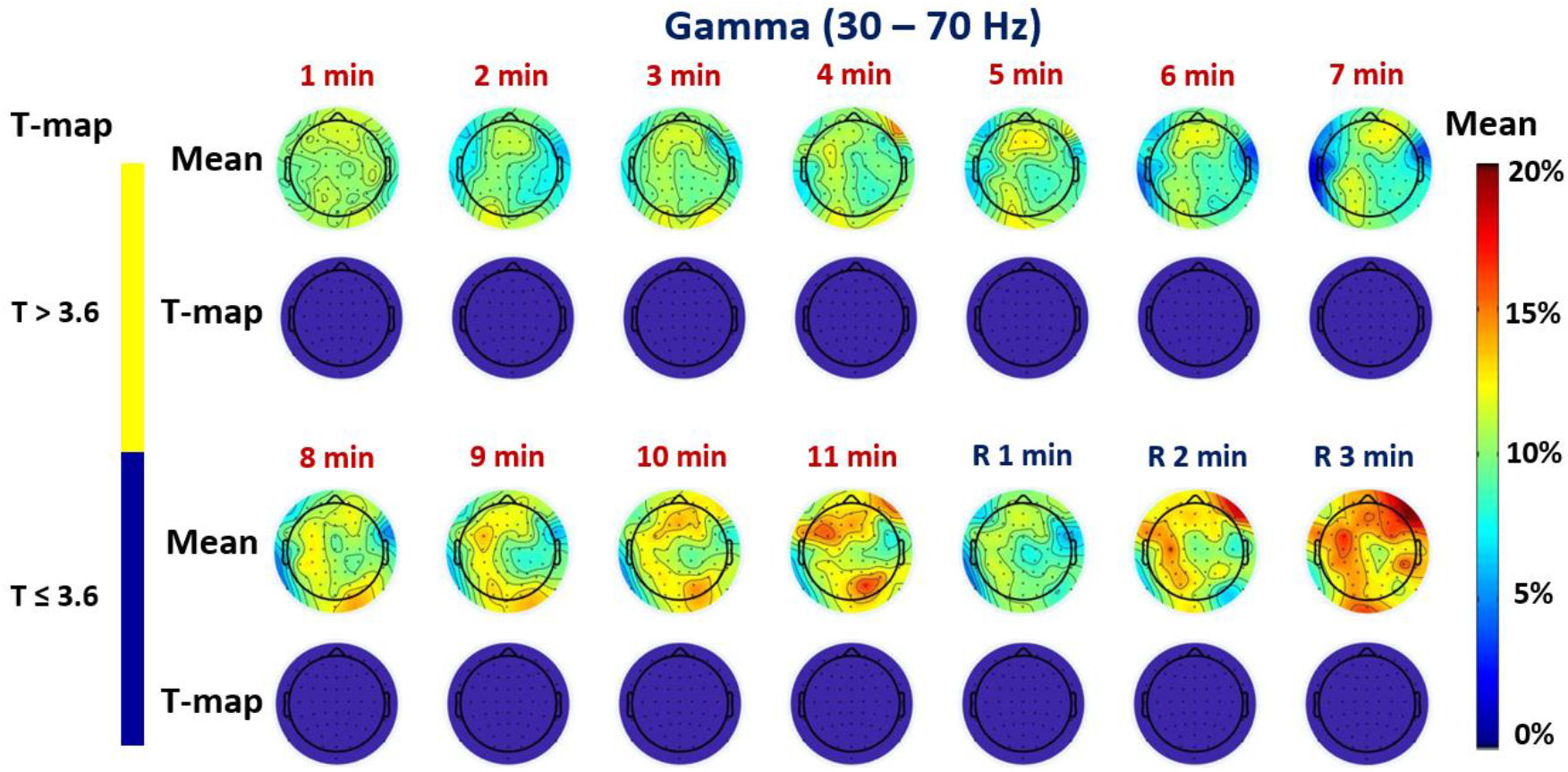
Time-resolved topographical maps of group-level differences in baseline-normalized EEG power density at gamma band between tPBM_L1064_ and sham experiments and corresponding statistical T-maps, during the 11 minutes of tPBM_L1064_/sham and 3 minutes of recovery period. The color bar on the right side represents mean difference in percentage increase of gamma power density between tPBM_L1064_ and sham stimulations. The color bar on the left represents the t-score cutoff for the Bonferroni-corrected p-value (p < 0.05, which is equivalent to T>3.6) of significant differences in power density between tPBM_L1064_ and sham stimulations.

## 4. Discussion

The findings provide the first detailed demonstration that tPBM_L1064_ significantly increases the strength of electrophysiological oscillations at the alpha and beta oscillations in different brain regions across the human scalp. Following a sham-controlled within-subjects design, we applied tPBM_L1064_ to the right forehead of 20 human subjects while recording the scalp EEG to investigate time-resolved, large-scale, electrophysiological changes across the entire scalp in response to tPBM_L1064_. Our multi-step analysis revealed several novel and intriguing findings, answered our research questions, proved our research hypothesis, and showed several limitations of this study.

### 4.1 tPBM_L1064_ modulates alpha and beta power

Figure 4 illustrates that the 11-min tPBM_L1064_ delivered to the right forehead of 20 human subjects (3 outliers excluded) resulted in up to 20% spatially broad increases of human alpha and beta band power. Figure 4(a) reveals that in the alpha band the scalp EEG oscillation powers increased progressively after the laser stimulation was initiated relative to the sham condition. Furthermore, the strongest increases in alpha oscillation power appeared during 8-11 min with a dominant front-to-back pattern bilaterally. Figure 4(b) shows similar increases for the beta band, but with a dominant pattern in posterior regions.

Together the figures reveal several key features: (1) laser-induced increases of EEG power density were <u>frequency-dependent</u> since the statistically significant increases occurred only at the alpha and beta bands, with very strong effects (maximum effect size 1.5); (2) laser-induced alterations were cumulatively <u>time-dependent</u> since the changes took place on a time scale of minutes for both alpha and beta bands, and they were largest during 8-11 min after laser onset; and (3) the alterations were <u>location-dependent</u>, based on the observations that significant increases in power density appeared to be front-to-back on the scalp 3 min after stimulation onset for the alpha and beta bands, with increases remaining front-to-back for the alpha band but only in the back for the beta band.

It is commonly regarded that an increase in alpha power alone implies a less attentive state and less arousal. It might be argued that the increase of alpha power seen in Fig. 4(a) could result from subjects’ sleepiness as they sat in the experiment for a while. If subjects get sleepy, alpha increases, but not both alpha and beta as produced by the laser as compared to the sham with the same temporal period. Therefore, the laser effect is not the same as simply lowering arousal level. This difference may be important for the cognitive benefit produced by the laser. It is also reported that alpha and beta frequencies of brain oscillations result from synchronous electrical activity of thalamo-cortical neurons, with alpha more characteristic of quiet awake states and beta of alert states (39, 40). More specifically, alpha waves are robustly modulated during cognitive processes and play an important role in the coalescence of brain activity (41). Beta oscillations are also relevant to cognitive processing but to a lesser degree than alpha waves (42, 43). Our results revealed that tPBM_L1064_ was able to create large effects on not only alpha but also beta oscillations. All of these observations led us to speculate that the laser-induced large effect in both alpha and beta power may be a mechanistic link to tPBM_L1064_-induced improvement and enhancement of human cognition that have been repeatedly reported in sham-controlled studies by Gonzalez-Lima et al. and others in recent years (12–15, 28). As supporting evidence for this speculation, many studies in neurofeedback research have observed improvement of cognitive performance while the upper alpha amplitude increased (44) by neurofeedback training (44–48).

### 4.2 tPBM_L1064_ holds potential for photobiomodulation of the human default mode network (DMN)

The DMN is a network of brain areas activated during the resting state; the DMN becomes suppressed once certain tasks are involved (49). Malfunction and disconnection of the DMN have been related to the cognitive dysfunctions seen in many psychiatric and neurological conditions, such as depression, anxiety and Alzheimer’s disease (49–51). The abnormal connectivity of the DMN in those conditions makes the brain unable to stay in a normal resting state (49). Increases of EEG power density in the anterior frontal regions indicate that tPBM_L1064_ may have the potential or ability to strengthen and stimulate neural functions of the anterior DMN. In particular, the T-maps of EEG alpha power during 8-11 min (Fig. 4(a)) illustrate that the scalp area covering (entirely or partially) the anterior DMN is significantly modulated by tPBM_L1064_. We note, however, that brain imaging modalities with higher spatial resolution than that of EEG will be needed to pinpoint the areas modulated by tPBM. The modulatory potential of tPBM_L1064_ on the DMN suggests a promising prospect for tPBM_L1064_ to promote DMN functions in healthy people and perhaps reverse DMN changes found in various psychiatric and neurological conditions such as depression, anxiety and Alzheimer’s disease.

### 4.3 Limitation of the study and future work

Since this was the first sham-controlled, time-resolved topographic study of EEG-tPBM_L1064_ in healthy humans, it had a few limitations that could be overcome in future investigations. First, to prevent the subjects from falling asleep during the EEG measurements, the subjects kept their eyes open during the measurements while wearing safety goggles with a layer of light-absorbing sheet over the edges for eye protection. However, in order to investigate tPBM_L1064_-induced effects thoroughly on alpha oscillations, it might be appropriate to conduct future experiments with the eyes closed, enabling detection of stronger alpha activity. Second, while the EEG data in this study were acquired using a conventional 10-10 system of scalp electrode placement, the EEG cap was positioned more posteriorly than the regular setup in order to have an adequate space for a 4-cm-diameter laser stimulation site (see Fig. 1(b)). The shift of the cap was approximately 2 cm up along the anterior-posterior midline (or the nasion to inion line), which would cause inconsistency between the conventional 10-10 system electrode sites versus the actual ones used in this study. This mismatch would prevent us from identifying individual scalp locations/regions that have significant EEG power responses to tPBM_L1064_ with high accuracy. A solution to this limitation may be to co-register the actual EEG electrodes with the conventional electrode placement using a 3-dimentional digitizer to measure several key anatomical landmarks, followed by coordinate transformation. Third, we speculated that the tPBM_L1064_-induced increase in both alpha and beta power density may be a mechanistic link to tPBM_L1064_-induced improvement and enhancement of human cognition that have been reported in sham-controlled studies in recent years (12–15, 28). This speculation needs to be further investigated and confirmed by combining EEG-tPBM_L1064_ measurements with tPBM_L1064_-enhanced human cognition tests in the future. Lastly, we cannot rule out that heating did not play a role in the electrophysiological changes observed here. Cortical temperature is known to change with neural activation by ~0.2 degrees Celsius (52). It is possible that the reverse effect – exogenous heat alters neural activity – could have contributed to the observed effects. On the other hand, we have previously shown that thermal stimulation of the forehead does not lead to the increased cortical CCO or hemodynamic changes with tPBM_L1064_ (53). In the future, a similar thermal stimulation study of EEG and computational simulations may be possible approaches to quantify or estimate tPBM_L1064_-induced vs. thermal effects on the EEG and thus be able to disambiguate the effects of photobiomodulation and tissue heating.

## 5. Conclusion

This is the first time-resolved topography report to demonstrate that tPBM with 1064-nm laser increases power of brain alpha and beta rhythm oscillations significantly and broadly across the scalp, in particular across bilateral anterior and posterior sites. The scalp EEG was utilized to measure sham-controlled, human electrophysiological rhythms before, during, and after 11-min tPBM_L1064_ delivered to the right forehead of a cohort of 20 normal human subjects. By computing oscillatory power in a time-resolved fashion, we found that tPBM_L1064_ produced time-varying statistically significant increases in EEG power in both the alpha and beta band. The electrodes of significant effects covered scalp regions with a front-to-back pattern, peaking 8 minutes after stimulation onset. Thus, tPBM modulated the synchronization of neural activity in the alpha and beta frequency range. Together with previous studies, this EEG study points to 1064-nm infrared light as a novel form of transcranial brain stimulation or neuromodulation by targeting cellular energy metabolism and associated changes in hemodynamics and electrophysiological activity of the brain.

## Acknowledgment

This work was supported in part by the National Institute of Mental Health/National Institutes of Health under the BRAIN Initiative (RF1MH114285). We also acknowledge the support in part from the STARS program by the University of Texas System. FGL was supported in part by the Oskar Fischer Project Fund.

